# Pupil diameter tracks statistical structure in the environment

**DOI:** 10.1101/770982

**Authors:** Caspar M. Schwiedrzik, Sandrin S. Sudmann

## Abstract

Pupil diameter determines how much light hits the retina, and thus, how much information is available for visual processing. This is regulated by a brainstem reflex pathway. Here, we investigate whether this pathway is under the control of internal models about the environment. If so, this would allow adjusting pupil dynamics to environmental statistics, and hence optimize information transmission. We manipulate environmental temporal statistics by presenting sequences of images that contain internal temporal structure to humans and macaque monkeys. We then measure whether the pupil tracks this structure not only at the rate that immediately arises from variations in luminance, but also at the rate of higher order statistics that are not available from luminance information alone. We find entrainment to environmental statistics in both species during the image sequences. Furthermore, pupil entrainment predicts later performance in an offline task that taps into the same internal models. Thus, the dynamics of the pupil are under control of internal models which adaptively match pupil diameter to the temporal structure of the environment, in line with an active sensing account.

## Introduction

Our sensory environment is richly structured in space and time: visual scenes and events unfold over time and present statistical regularities [1]. Our sensory systems can extract these statistics to form internal models that in turn allow optimizing perceptual processing. Such internal models can also be used to guide sampling of relevant information through appropriate motor actions [2]. This is captured by the notion of “active sensing” [3]: for example, we can use visual information to plan sequences of consecutive saccades that target the most relevant locations in a scene, thus providing the visual system with bouts of information, e.g., for object recognition. A critical component of active sensing in vision is how much light and therefore how much information may hit the retina. This is regulated by the reflexive adjustment of pupil diameter. Here, we investigate whether this adjustment is fully automatic or under the control of flexible internal models.

When light hits the retina, the pupil transiently constricts to limit light influx. This pupillary light response (PLR) is controlled by a parasympathetic brainstem circuit in which luminance information from the retina is relayed via the pretectal olivary nucleus to the Edinger-Westphal nucleus, which in turn signals the pupillary sphincter muscle to contract. In addition, a sympathetic pathway adjusts pupil diameter to background illumination [4]. By doing so, pupil size modulates activity in visual cortex which scales with the amount of light passed [5]. Pupil diameter affects acuity and sensitivity of visual processing [6]: smaller pupils sharpen the image and increase depth of field, while larger pupils allow more light to hit the retina and thus increase the signal-to-noise ratio and the field of view. This suggests that visual processing can be optimized by adjusting pupil diameter to environmental conditions to maximize information transmission, in line with an active sensing account [7, 8]. Some environmental statistics are not immediately available from light intensity alone and require more complex computations. Indeed, there is evidence that the PLR is not a fully automated reflex but under some degree of cortical or collicular [9] control. For example, the PLR is modulated by chromatic isoluminant stimuli, which cannot be the result of isolated subcortical processes that do not have access to chromaticity [10]. Here, we ask whether pupil diameter adjusts to higher-order *temporal* statistics in the visual input, i.e., whether the pupil can entrain to temporal structure that is not a direct consequence of light variations but needs to be derived from higher-order environmental regularities. By adjusting its temporal dynamics to the environment, the pupil could act as an adaptative filter that optimizes the signal-to-noise ratio at a given, environmentally relevant frequency. If so, this would constitute a sophisticated mechanism of active sensing whereby internal models act on the very onset of visual processing.

We artificially induced temporal statistics by presenting sequences of achromatic, luminance-equalized images at a fixed temporal rate (2 Hz) while manipulating their internal order: we grouped individual images into pairs, such that the first image in a pair predicted the identity of the second image (Figure 1A). This way, we could separate the luminance-induced PLR occurring at the 2 Hz image-rate from any modulation of pupil diameter at the rate of pairs at half that frequency, 1 Hz. Because only the statistical structure of the stimulus streams but not the variations in light intensity can possibly drive a 1 Hz response under these conditions, a modulation of pupil diameter at 1 Hz would speak for sensitivity of the responsible subcortical pathways to higher-order environmental statistics. To test whether this effect reflects a common sensorimotor strategy across species, we conducted experiments in humans and macaque monkeys. While the brainstem circuit for the PLR is preserved between these two species [11], there are systematic differences in pupil dynamics between monkeys and humans [12], and it is unclear whether the pupil is under the same amount of cortical/cognitive control.

**Figure 1.**
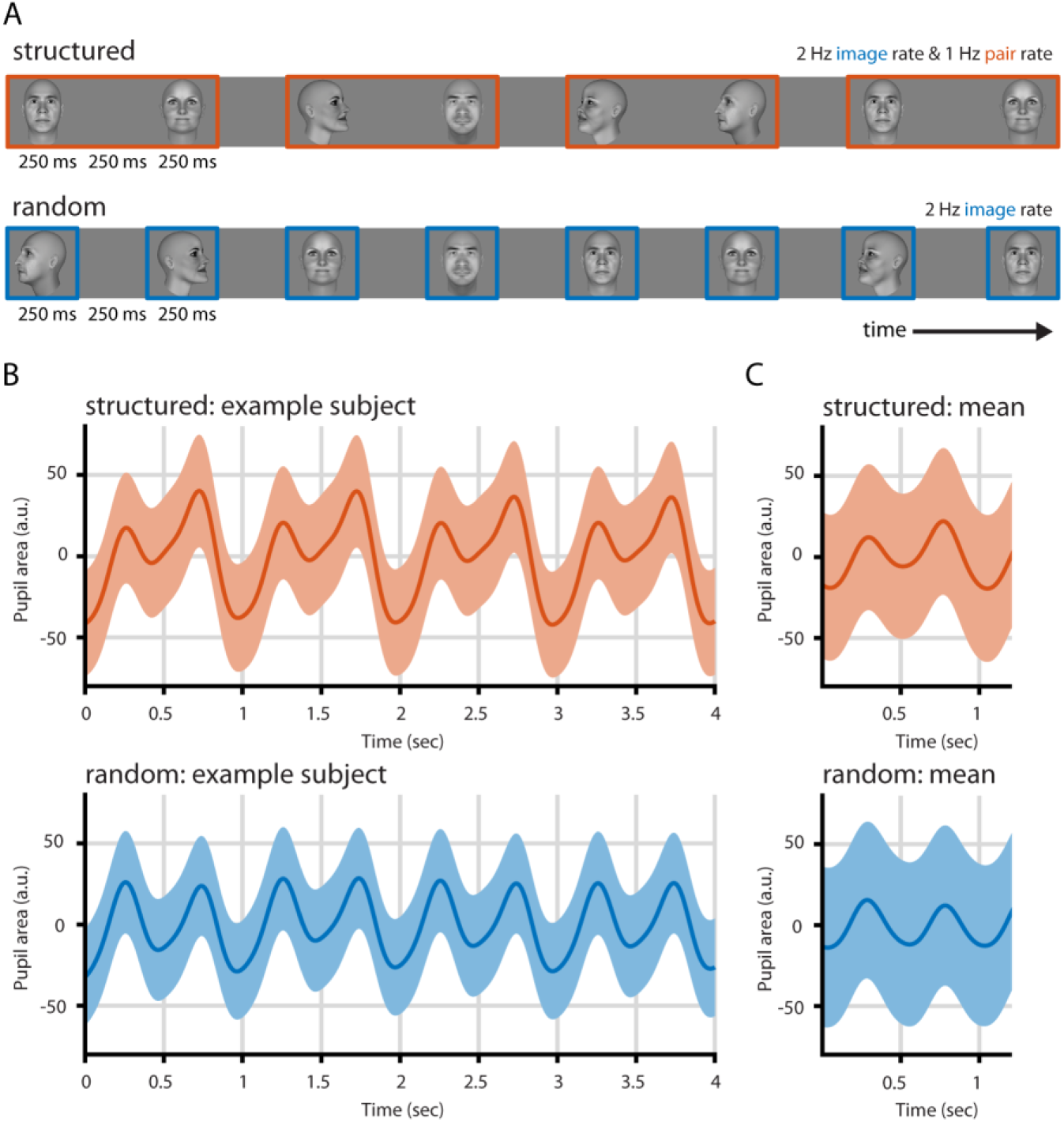
Paradigm and pupil entrainment. **(A)** We presented sequences of faces in structured or random order. In the random condition, faces were shown in random order at 2 Hz, the image rate. In the structured condition, faces were shown at the same rate (2 Hz). To induce statistical structure, images were grouped into pairs, such that one particular image always followed on other particular image. This gives rise to the pair rate at 1 Hz. **(B)** Pupil entrainment was evident at the single subject level (here: mean over 83 trials). The structured condition (orange) shows a clear modulation at the 2 Hz image rate and slower dynamics, including the pair rate at 1 Hz. The predominant frequency in the random condition (blue) is the image rate (2 Hz). **(C)** Pupil entrainment at 1 and 2 Hz is also clearly evident when averaged across all runs and subjects. In B and C, grey vertical lines indicate image onsets, shading represents the SEM, and data are lightly detrended.

## Results

To test whether pupil diameter is sensitive to temporal environmental statistics, we presented long sequences (2 min or longer) of computer-generated, luminance-equated face stimuli varying in facial identity and head orientation against a grey background (Figure 1A, Supplemental Figure S1). Each face was shown for 250 ms, followed by a 250 ms gap. Such alternation in light intensity should induce a pupillary response at 2 Hz, the image rate. This is well within the range of pupillary constriction dynamics [13]. In the random condition, the images were presented at a fixed rate but in random order. In the structured condition, we presented the same images at the same rate, but we arranged them such that they were systematically grouped into pairs, i.e., the transitional probabilities between specific images were fixed at 100%, while the transitional probabilities between pairs were minimized. Thus, in addition to the individual images, this pairing gave rise to temporally coherent units at half the image rate. If this statistical structure was extracted, it should be reflected by a modulation of the pupil at 1 Hz, the pair rate. To draw attention to the stimuli, but not explicitly to their temporal structure [14], human subjects were instructed to perform a 1-back task during both conditions in which they had to detect infrequent repetitions of identical faces. Performance in the task was above chance (random: mean accuracy 59.97%, *t*(29)=4.786, *p*<0.001, *g*=0.851; structured: mean accuracy 59.72%, *t*(29)=3.147, *p*=0.004, *g*=0.560) and did not differ between conditions (mean difference 0.25%, *t*(29)=0.095, *p*=0.925, *g*=0.017).

We first present the human data. Entrainment of the PLR at the 2 Hz image rate was clearly visible in the pupil signal on the single subject level (Figure 1B). Spectral analyses of the continuous data showed a distinct peak in spectral power at the image rate (2 Hz) in both the structured (*t*(29)=15.018, *p*<0.001, *g*=2.314) and the random (*t*(29)=13.494, *p*<0.001, *g*=2.134) condition compared to the four surrounding frequency bins (Figure 2). In addition, we find a strong response at 1 Hz, the pair rate, but only in the structured condition (interaction condition × frequency, *F*(1,29)=6.114, *p*=0.019, *η*^*2*^=0.174; structured vs. random, mean difference 1.493 dB, *t*(29)=3.451, *p*=0.002, *g*=0.449). In fact, there was no distinct peak at 1 Hz in the random condition compared to the surrounding bins (*t*(29)=0.519, *p*=0.607, *g*=0.038). A complementary analysis of pupillary phase revealed significantly stronger phase locking at 1 Hz in the structured than in the random condition (mean difference 0.151, *t*(29)=5.326, *p*<0.001, *g*=1.447). Control analyses showed that the 1 Hz peak was not evident in the concurrently recorded eye movement signal in either condition (Supplemental Figure S2), and that there were no correlations between 1 Hz spectral power in the pupil and the eye movement signal (horizontal: *r*=-0.264, *p*=0.159; vertical: *r*=0.117, *p*=0.539). This rules out that pupil entrainment at this frequency was an artifact arising from other eye movements and blinks that may occur at a similar rate. Thus, the human pupil can track environmental statistics that are not directly evident in the light pattern that hits the retina.

**Figure 2.**
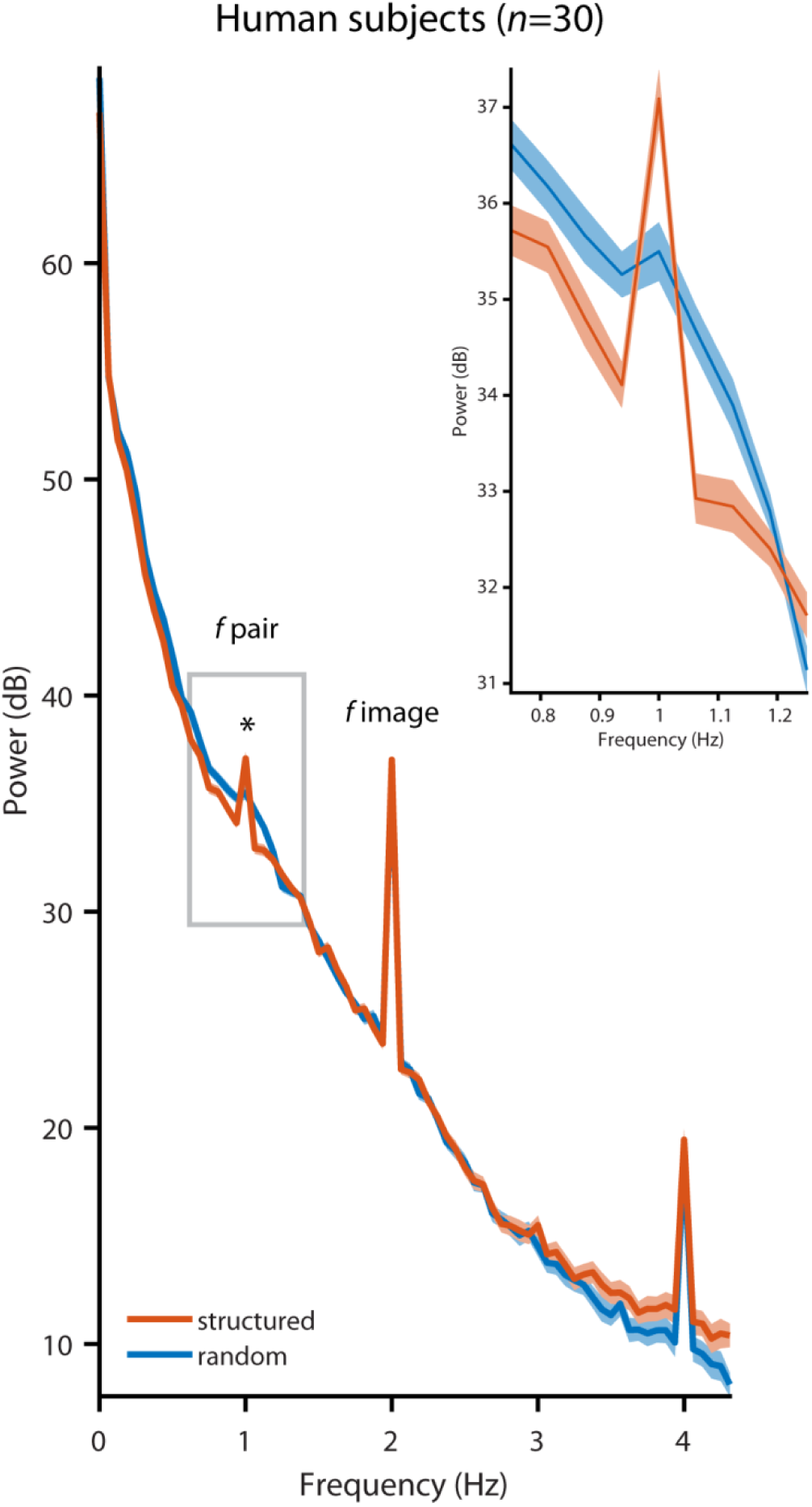
Pupil diameter in human subjects (*n*=30) followed the image rate at 2 Hz in the random (blue, *t*(29)=13.494, *p*<0.001, *g*=2.140) and the structured (orange, *t*(29)=15.018, *p*<0.001, *g*=2.314) condition (compared to the mean of the four surrounding frequencies), but the pair rate at 1 Hz was only evident in the structured condition (structured vs. random, *t*(29)=3.451, *p*=0.002, *g*=0.449). The inset shows a zoomed version of the power spectrum around 1 Hz. Peaks at 4 Hz likely reflect harmonics. Shading represents the SEM across subjects, corrected for inter-subject variability [16].

To determine which aspects of pupillary movement contribute to the dynamics at 1 Hz in the structured condition, we examined pupil dilations and constrictions, respectively, time-locked to the onset of the pairs. Pupil dilation can be elicited by surprising stimuli under constant illumination [15]. In our paradigm, once the statistics of the structured stream have been acquired, they may render stimuli more or less predictable/surprising. Specifically, the second stimulus in a pair is entirely predictable from stimulus 1. In contrast, the first stimulus in a pair may be surprising as it cannot be predicted from its predecessor. Hence, transient pupil dilations to every first stimulus reflecting surprise could contribute to spectral power at 1 Hz. We find, however, that pupil dilation following the first stimulus was smaller than following the second stimulus (in the structured condition only, interaction condition × time point, *F*(1,29)=18.526, p<0.001, *η*^*2*^=0.390). This suggests that in our paradigm, pupil dilation at 1 Hz reflects statistical structure of the input stream but is not (solely) driven by surprise.

In addition, Figure 1B and C show how pupil constriction may contribute to the 1 Hz response in the structured condition: pupil constriction between images is suppressed within a pair, while pupil constriction between pairs appears particularly pronounced. Indeed, across subjects, mean pupil constriction was significantly smaller within pairs than between pairs only during the structured condition (interaction condition × time point, *F*(1,29)=14.043, *p*<0.001, *η*^*2*^=0.326). This suggests that the constrictive PLR *within* pairs is diminished as if not to interrupt information transmission within an environmentally coherent unit, and augmented *between* pairs, possibly to segment the continuous input stream at the pair rate. Taken together, this shows that both dilation and constriction dynamics contribute to the 1 Hz response in the structured condition, with the net effect of an overall wider pupil within a pair than between pairs.

We ran the same experiments with macaque monkeys, albeit with a reduced number of stimulus pairs (3 instead of 9). Instead of performing the 1-back task, monkeys passively viewed the stimulus sequences and were rewarded for maintaining fixation. Like in humans, we find in both monkeys a PLR at 2 Hz in the structured and the random condition, but a response at 1 Hz only when there is statistical structure (Figure 3; condition × frequency interaction, Monkey P: *F*(1,31)=69.610, *p*<0.001, *η*^*2*^=0.692; monkey C: *F*(1,41)=18.216, *p*=0.001, *η*^*2*^=0.308). The strength of the modulation at 1 Hz, relative to the surrounding frequencies, in both monkeys (mean difference 4.97 dB monkey P, 1.25 dB monkey C) fell well into the range we observed in humans (min 0.15 dB, max 6.32 dB). Bayesian statistics showed that the probability that the difference for human subjects would be more extreme than in our monkey subjects was only 5.12% (monkey P) and 25.51% (monkey C), suggesting that monkeys and humans showed similar pupillary dynamics. As in humans, both monkeys also showed stronger phase locking at 1 Hz in the structured than in the random condition when we analyzed pupillary phase (monkey P: *t*(31)=13.361, *p*<0.001, *g*=4.540; monkey C: *t*(41)=5.582, *p*<0.001, *g*=1.753). There were no significant differences between conditions in other eye movements (Supplemental Figure S3). Thus, the cognitive modulation of pupil diameter based on environmental statistics we found in humans is shared with macaques, whose evolutionary lineage split from ours some 25 million years ago [17].

**Figure 3.**
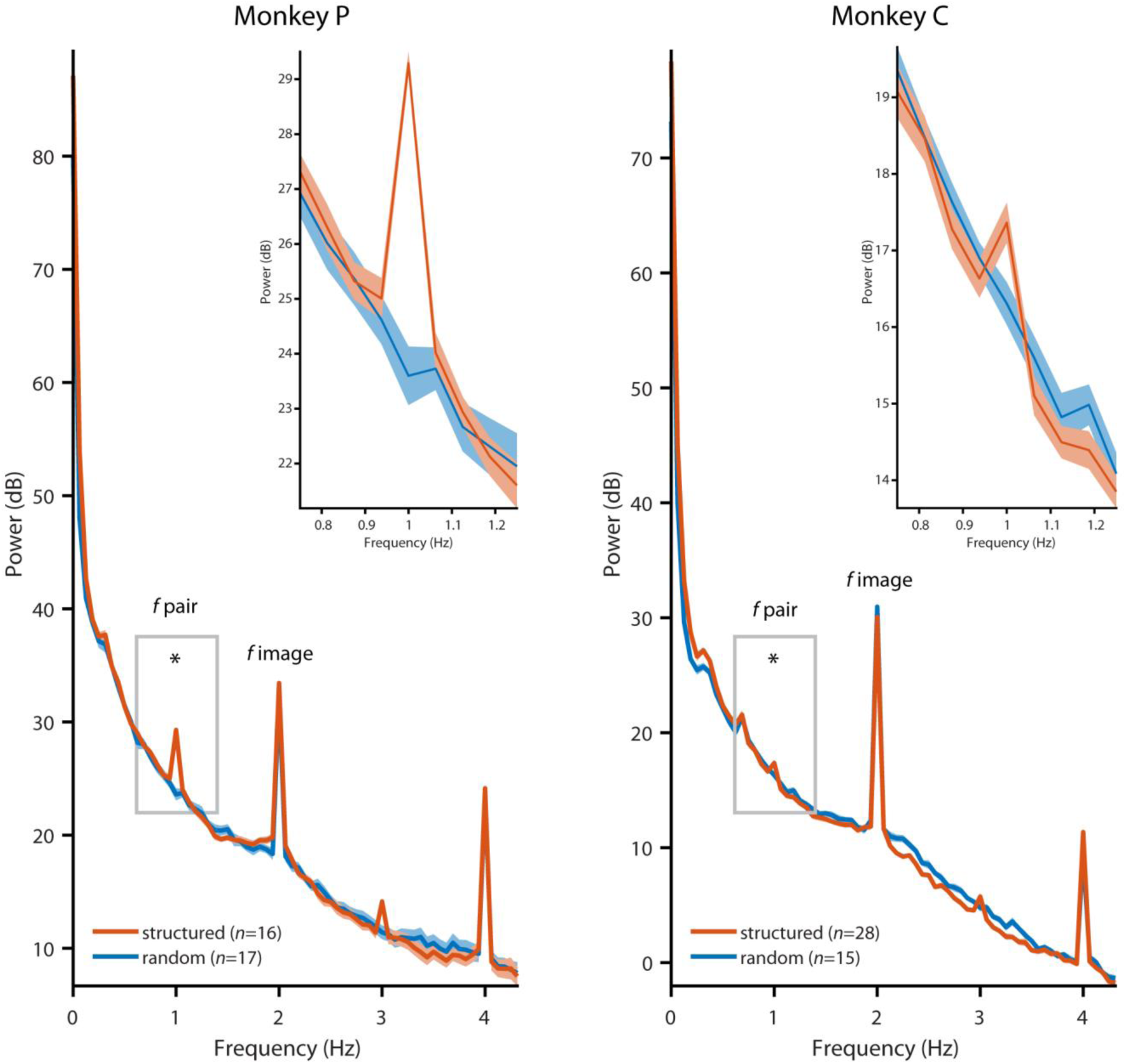
Pupil diameter in monkeys followed the image rate at 2 Hz in the random (blue) and the structured (orange) condition, but the pair rate at 1 Hz was only evident in the structured condition (structured vs. random, monkey P:, *t*(31)=10.37, *p*<0.001, *g*=3.524; monkey C: *t*(41)=2.327, *p*=0.025, *g*=0.731). The inset show a zoomed in version of the power spectrum around 1 Hz. Peaks at 3 and 4 Hz likely reflect harmonics of the 1 and 2 Hz response, respectively. Shading represents the SEM across sessions.

If pupil diameter entrains to environmental statistics to optimize the transmission of visual information about pairs, this may have perceptual consequences. We thus tested in human subjects whether pupil entrainment predicted perceptual benefits during a task in which the statistics were directly relevant for task performance. Specifically, after watching the random and subsequently the structured sequences, subjects performed a new task in which they had to detect a face stimulus in a rapid serial visual presentation (RSVP) sequence of other faces (Figure 4A). In a majority of the trials, the target face was immediately preceded by the face with which it was paired during the structured exposure phase (the predictor) to induce a priming effect. For comparison, we created two additional test conditions that violated the previously exposed structure: the ‘foil’ condition, in which we replaced the predictor with a facial identity or head orientation that had been shown during exposure but that had been paired with a different target; and the ‘novel’ condition, in which we replaced the predictor with a novel facial identity. We hypothesized that if subjects learned the order and identity of the face pairs during the structured exposure phase, then violating learned associations during the RSVP should lead to lower accuracy and slower reaction times during target detection relative to detecting a target that appears in the known configuration.

**Figure 4.**
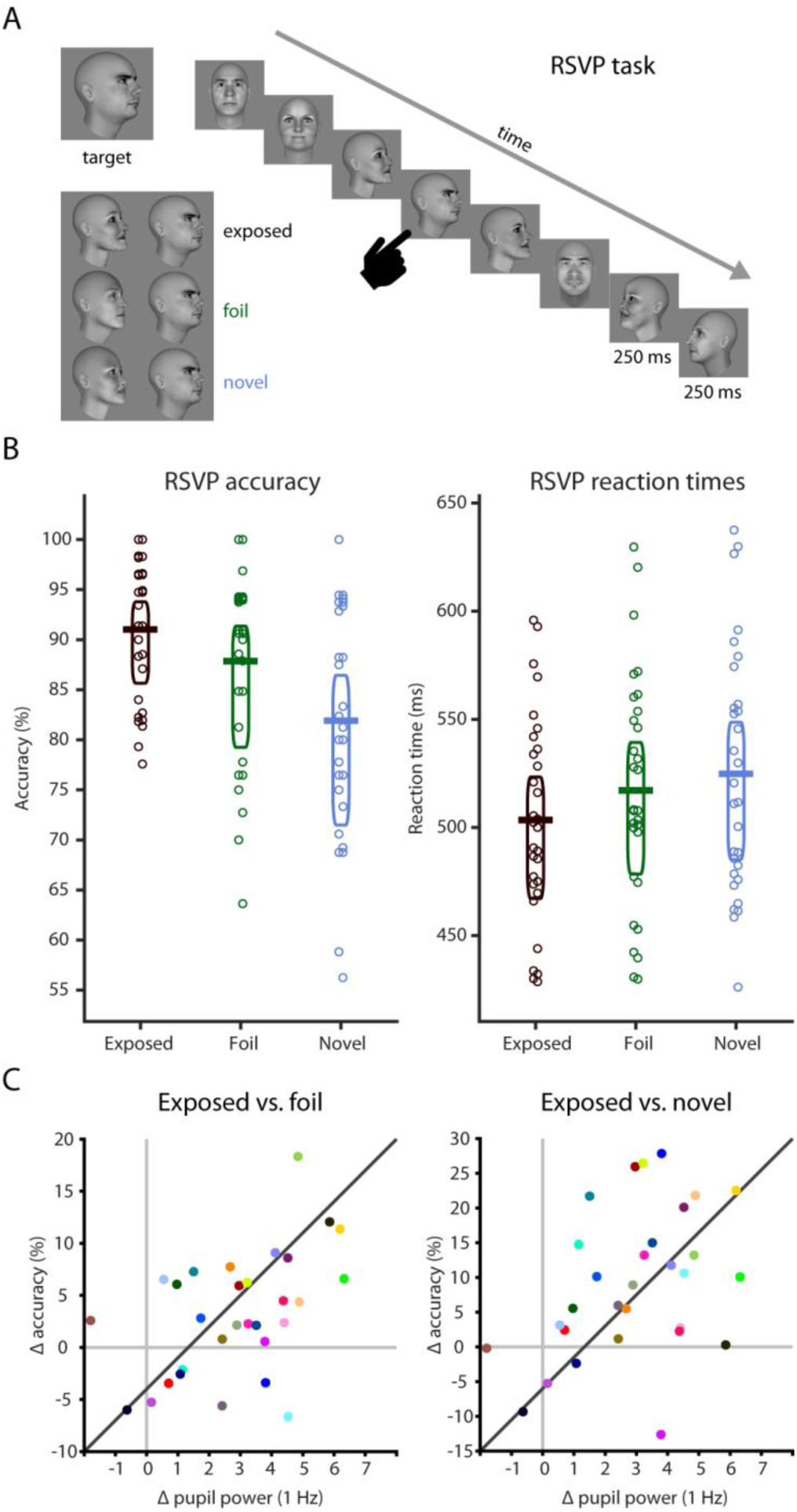
Offline learning test and relation to pupil entrainment. **(A)** On each trial of the RSVP task, subjects had to detect a target face. Targets were embedded into a stream of faces (250 ms on, 250 ms off). The face immediately preceding the target could be the previously exposed predictor (“exposed” condition, black), an identity or a head orientation that had been paired with another face during the exposure phase (“foil” condition, green), or a completely novel identity (“novel condition”, blue). **(B)** Human subjects were more accurate (exposed vs. foil, *t*(29)=2.928, *p*=0.007, *g*=0.376; exposed vs. novel, *t*(29)=4.715, *p*<0.001, *g*=0.962) and faster (exposed vs. foil, *t*(29)=-4.522, *p*<0.001, *g*=-0.266; exposed vs. novel, *t*(29)=-5.464, *p*<0.001, *g*=-0.403) in detecting the target when it was preceded by the previously exposed predictor than in the foil or the novel conditions. Horizontal bars indicate the mean, boxes the 95% Bayes-bootstrapped high density interval, circles the individual subjects’ data points (*n*=30). **(C)** Pupil entrainment at the 1 Hz pair rate during the exposure phase (normalized against the mean of the four surrounding frequencies) predicted offline learning effects: the stronger the entrainment at 1 Hz, the larger the accuracy benefit in the exposed over the foil (*r*=0.481, *p*=0.007) and the novel (*r*=0.402, *p*=0.027) condition.

Indeed, subjects were significantly faster (exposed vs. foil, mean difference -13.692 ms, *t*(29)=-4.522, *p*<0.001, *g*=-0.266; exposed vs. novel, mean difference -21.344 ms, *t*(29)=-5.464, *p*<0.001, *g*=-0.403) and more accurate (exposed vs. foil, mean difference 3.17%, *t*(29)=2.928, *p*=0.007, *g*=0.376; exposed vs. novel, mean difference 9.11%, *t*(29)=4.715, *p*<0.001, *g*=0.962) in detecting targets in the exposed than in the foil and in the novel condition (Figure 4B), evidencing that they indeed extracted and retained the statistics of the face sequences from the exposure phase. We then correlated the performance differences in accuracy from the RSVP task with the degree of pupil entrainment at the 1 Hz pair rate during the preceding exposure phase (Figure 4C). We find that the stronger the pupil entrainment during exposure, the larger the accuracy benefit of the exposed over the test conditions (exposed vs. foil *r*=0.481, *p*=0.007; exposed vs. novel *r*=0.402, *p*=0.027) in the RSVP task. The same held true when controlling for any correlation between 1 Hz spectral power in the random condition and accuracy (partial correlation; exposed vs. foil *r*=0.450, *p*=0.014; exposed vs. novel *r*=0.412, *p*=0.026). Hence, pupil entrainment during exposure predicts subsequent detection performance, suggesting that they are based on the same internal models.

## Discussion

We find that the peripheral oculomotor system is involved in actively tracking environmental statistics, in line with an active sensing account. The modulation of pupil diameter is conserved between monkeys and humans. The latter is not a given, as other modulations of pupil diameter show systematic differences between the two species that point to differences in the underlying anatomy and/or neural pathways involved [12]. In both species, the pupil entrains to higher-order structure in the visual input, which leads to an adaptive correlation between visual ecology and response dynamics: the visuomotor system matches the dilation-constriction dynamics of the pupil to the temporal structure of the environment. This involves an increase in pupil dilation for predictable stimuli, and a reduction of pupil constriction within a pair. The result is an overall wider pupil during the presentation of environmentally coherent units, which may maximize information transmission for downstream visual processing. In addition, we find an augmented PLR between pairs, akin to an active segmentation of the continuous input stream at event boundaries between coherent units. Taken together, this resembles an adaptive filter in the time domain that optimizes the signal-to-noise ratio at ecologically relevant frequencies.

Our findings imply that pupil-controlling pathways in the brainstem are under control of (or at least influenced by) brain areas that can extract environmental statistics and translate them into useful models for active sampling already at the very first stage of visual interaction with the outside world. Previous research has shown that pupil diameter is modulated by other factors than light influx which likely arise from cortical computations, e.g., attention [18, 19], motion coherence [10] and even speech [20]. One possible source of pupil diameter modulation is extrastriate visual cortex, where the same paradigm we used here affects neural processing of facial information [21] and where lesions affect pupil diameter modulation by other visual features like color [22]. Statistical structure also entrains brain areas beyond visual cortex, including motor and premotor cortex [23]. Here, stimulation, e.g., of the frontal eye fields, induces pupil size changes in macaque monkeys [24, 25]. Alternatively or additionally, modulation of the pupil may be mediated by the superior colliculus [9] or the locus coeruleus [26], whose activity is also known to covary with pupil diameter.

Finally, on a technical note, pupil entrainment presents itself as an easy to obtain, high signal-to-noise behavioral readout of learning models from the environment, possibly permitting assessment of sensitivity to statistical structure across levels of complexity between individuals, stages of development, or species.

## Acknowledgements

We would like to thank I. Kagan for support, J. Fischer and L. Melloni for comments on the manuscript, L. Burchardt, D. Bertazzi Lazzarini, and R. Brockhausen for technical support, T. Becker for veterinary care, and A.-L. Beyer for help with data acquisition. This research was supported by an Emmy Noether grant from the German Research Foundation (SCHW1683/2-1) to C.M.S..

## Author contributions

C.M.S. conceived the experiments. S.S.S. performed experiments. C.M.S. and S.S.S. performed data analysis and wrote the manuscript.

## Methods

### Subjects

33 healthy human volunteers (20 female, 2 left handed, mean age 25.45 yrs, SD 3.46 yrs) participated in this study. All subjects had normal or corrected-to-normal vision, reported no history of neurological or psychiatric disease, and gave written informed consent before participation. No sample size estimate was performed, but sample size was selected based on previous studies. Three subjects had to be excluded from data analysis because they did not complete the study or failed to follow instructions (final *n*=30, 19 female, 2 left handed, mean age 25.07 yrs, SD 2.90 yrs). All procedures with human subjects were approved by the Ethics Committee of the University Medical Center Göttingen (protocol number 29/8/17). Subjects received monetary compensation for their participation.

In addition, two adult male rhesus monkeys (*Macaca mulatta*) participated in the study (age 7 yrs (monkey P) and 13 yrs (monkey C), weight 7.3 kg (monkey P) and 8.4 kg (monkey C) at time of testing). Sample size matched that of earlier studies [12]. Both animals had previously been implanted with cranial head-posts under general anesthesia and aseptic conditions, for participation in neurophysiological experiments. The surgical procedures and purpose of these implants were previously described in detail [21, 27]. Animals were extensively trained with positive reinforcement [28] to enter into and stay seated in a primate chair, and to have their head position stabilized via the head-post implant. This allows implant cleaning, precise recordings of gaze and neurophysiological recordings while the animals work on cognitive tasks. Here, we made opportunistic use of these situations to record eye movement and pupil dilation data. The experimental procedures were approved by the responsible regional government office (Niedersächsisches Landesamt für Verbraucherschutz und Lebensmittelsicherheit - LAVES). The animals were pair- or group-housed in accordance with all applicable German and European regulations. The facility provides the animals with an enriched environment (including a multitude of toys and wooden structures), natural as well as artificial light and access to outdoor space, exceeding the size requirements of European regulations. The animals’ psychological and veterinary welfare was monitored daily by veterinarians, animal facility staff and scientists.

### Stimuli and tasks

We used a set of images depicting human faces from a previous study [21]. In brief, we generated 36 3-dimensional human faces with a neutral expression and no hair in FaceGen (v3.5.3, Singular Inversions). Images were converted to black and white and luminance normalized using SHINE [29]. For this study, we selected 18 unique images from the full set, each showing a different face (Supplemental Figure S1). For humans, images (10 × 10 dva) were presented foveally on an LCD monitor (ViewPixx EEG, refresh rate 120 Hz, resolution 1920 × 1080 pixel, viewing distance 68 cm) in a darkened, sound-attenuating booth (Desone Modular Acoustics). For monkeys, images (40 × 40 dva) were presented using a projector (Barco F22 WUXGA, refresh rate 60 Hz, resolution 1920 × 1080 pixel, viewing distance 52 cm) in a darkened training setup. Stimulus delivery and response collection for human subjects were controlled using Presentation (v19, Neurobehavioral Systems); visual stimulation and reward for monkey subjects were controlled using MWorks (https://mworks.github.io/).

During the exposure phase, subjects viewed the faces in temporal sequence (Figure 1). In the random condition, images were presented in random order for 250 ms each, with a 250 ms inter-stimulus interval (ISI). Human subjects were exposed to 1188 images in six blocks of 198 images within a single session. Monkey subjects were exposed to an average of 6000 images per session and completed 38 (monkey P) and 46 (monkey C) blocks of 1200 images in which fixation accuracy was >=85%, respectively. Monkey P completed 19 blocks in the random and 19 blocks in the structured condition (see below); monkey C completed 17 blocks in the random and 29 blocks in the structured condition. Human subjects were instructed to fixate on a blue fixation dot. To assure that subjects were paying attention to the images, they performed a 1-back repetition detection task, i.e., they had to report an infrequent (18/198 per block) immediate repetition of an identical image by means of a button press on a standard keyboard. Monkeys passively viewed the stimuli but were rewarded with juice or water if they continuously fixated on a red, centrally presented fixation dot for 2-4 s. Effective fixation accuracy was <2 dva (median 95%, MAD 1.96).

Stimuli and timing were identical in the structured condition, but unbeknownst to the subjects, images were now joined into pairs (Supplemental Figure 1). Pairs were arranged such that one identity-view combination would uniquely predict one other identity-view combination, while assuring that head orientation was fully balanced across pairs (e.g., 3 different identities at 0 degrees head orientation were paired with 3 different identities at 0, 60 and 300 degrees head orientation, respectively). To induce statistical structure, the sequence of pairs was arranged such that transition probabilities within pairs (i.e., between stimuli) were 100%, while transition probabilities between pairs (i.e., between trials) were at minimum and balanced across pairs. Human subjects were exposed to nine pairs, monkey subjects to three. Subjects performed the same tasks as in the random condition. Because reward for monkey subjects was solely delivered on the basis of fixation performance, there was no systematic relationship between the occurrence of a pair and reward.

In addition, we tested whether human subjects retained the statistical structure they had been exposed to during the structured condition in a subsequent offline test (Figure 4A). Specifically, subjects had to detect a target face in a rapid serial visual presentation (RSVP) stream of face images by means of a speeded button press [14]. On each trial, we first presented the target image above the central fixation dot. The target was one of the second images from the 9 face pairs. The subjects could then initiate the RSVP by a button press. The RSVP consisted of 18 face images presented at fixation and with the same timing as during the exposure phase (250 ms stimulus duration, 250 ms ISI). The target image could not appear as the first or last image in the sequence. As in the structured exposure condition, and again unbeknownst to the subjects, all images in the RSVP were presented as pairs. This served to assess whether the subjects had acquired and retained the statistics of the structured exposure phase. If so, we hypothesized that they could predict the occurrence of the second image in a pair once they saw the corresponding first image, and that this should facilitate face detection [21]. To test this hypothesis, we created two test conditions that violated the exposed associations: in the ‘foil’ condition, we replaced the first image in a pair that immediately preceded the target face during the RSVP either with a facial identity that had been shown during exposure but that had been paired with a different target, or with an image that showed the previously paired identity but with a different head orientation. In the ‘novel’ condition, we replaced the first image in the pair by a novel facial identity (but preserving the head orientation from the exposure phase). If subjects had acquired image-specific predictions about the order and identity of the face pairs during the structured exposure phase, then violating these expectations during the RSVP should lead to lower accuracy and slower reaction times during target detection relative to detecting a target that appears in the known configuration. Subjects completed 63 trials with pairs in the previously exposed configuration (seven per pair), 36 trials in the ‘foil’ condition, and 18 trials in the ‘novel’ condition, a total of 117 trials. We retained a ratio of 1.2 trained over test conditions as to not overwrite expectations stemming from the preceding structured exposure phase.

### Pupil and gaze measurements

During all experiments, we continuously acquired pupil and gaze measurements using a high-speed, video-based eye tracker (SR Research Eyelink 1000+). Data were sampled at 1000 Hz from both eyes.

### Data analysis

#### Pupil preprocessing

All data analyses were carried out in Matlab (R2017b, The Mathworks, Inc.). We preprocessed pupil area data by first linearly interpolating blinks, otherwise missing values, as well as outliers exceeding 3.5x the median absolute deviation (MAD) over the entire recording per eye per subject. Data were then lowpass filtered at 5 Hz using a onepass-zerophase Kaiser-windowed sinc finite impulse response (FIR) filter (filter order 1812, transition width 2.0 Hz, pass band 0-4.0 Hz, stop band 6.0-500 Hz, maximal pass band deviation 0.0010 (0.10%), stop band attenuation -60 dB) [30]. Subsequently, data were detrended per block and the mean per block as well as an average over the baseline of 2 s before the beginning of stimulation in each block were subtracted. Data were then averaged between eyes and finally downsampled to 500 Hz (keeping every second sample). We also corrected pupil data for gaze position [31] before filtering, but this did not affect the pattern of results. We thus report uncorrected pupil data.

#### Pupil spectral power

For analyses of spectral power, we performed a Fourier transform per block using Welch’s method [32], as implemented in the Matlab function *pwelch.m*. We used a Hanning window of 16866 points (1/3 of the block length), 5622 overlapping points (1/3 of the window length), and 8000 discrete DFT points to obtain a spectral resolution of 0.0625 Hz. The same preprocessing and analysis steps were carried out for the horizontal and vertical eye position signals. Before statistical analyses, blocks in which the preprocessed pupil signal exhibited large remaining artifacts or in which the power spectra were distributed abnormally were excluded (13.3% of human data, 9.5% of monkey data). The remaining power spectra were converted to dB by taking the decadic logarithm and multiplying by 10. In humans, the block wise power spectra were averaged per subject and compared statistically across participants. In monkeys, statistical analyses were done comparing blocks per animal. We specifically focused on the pair frequency at 1 Hz and the image frequency at 2 Hz. To statistically determine the existence of spectral peaks in the power spectra, we compared spectral power at 1 Hz and 2 Hz, respectively, to the average of the four surrounding frequency bins (two above, two below), by means of a paired *t*-test. Figures S2 and S3 show the respective mean differences and corresponding 95% Bayes-bootstrapped (1000 samples) high density intervals and were plotted using the Robust Statistical Toolbox (https://github.com/CPernet/Robust_Statistical_Toolbox) in Matlab. To assess whether there were statistically significant differences between the random and the structured condition and whether they were specific to the pair/image frequency, we carried out a repeated measures analysis of variance (rmANOVA) with frequency and condition as within subjects factors in humans and a mixed ANOVA with frequency as within blocks and condition as between blocks factor in monkeys. In addition, we performed paired *t*-tests between conditions at the pair and image frequency, respectively, in humans, and independent samples *t*-tests in monkeys. To assess whether pupil dilation differed after the presentation of the first and the presentation of the second stimulus in a pair, we compared average pupil dilation between 250 and 350 ms after the onset of the first and second stimulus, respectively, during the random and the structured condition, respectively, in humans using a rmANOVA. To assess whether pupil constriction differed within and between pairs, we compared the average pupil constriction between 500 and 600 ms after pair onset (within) to the average pupil constriction between 1000 and 1100 ms after pair onset (between) during the random and the structured condition, respectively, in humans using a rmANOVA. To compare the modulation of pupil dynamics at 1 Hz between species, we used a two-tailed, Bayesian Standardized Difference Test (https://homepages.abdn.ac.uk/j.crawford/pages/dept/psychom.htm) [33] that allows comparing individual subjects, in our case individual monkeys, to a group of subjects, in our case the human observers, and estimates the probability that a more extreme score than the one observed in the respective individual subject is found in the group.

#### Pupil phase consistency

We also assessed whether there was consistent phase locking of the pupil signal to the pairs at 1 Hz. To this end, we cut the continuous data into pseudo-trials of 32 stimuli (starting with every second stimulus) after the filtering preprocessing step, baseline-corrected each pseudo-trial by subtracting an average over the 2 s pre-block baseline, and averaged the signal between eyes. We then computed a trial-by-trial Fourier transform using discrete prolate spheroidal sequences as tapers with the same spectral resolution as for the analyses of spectral power (0.0625 Hz), using the Matlab toolbox Fieldtrip (v20170327; http://www.fieldtriptoolbox.org/) [34]. The resulting complex spectra were then used to calculate inter-trial phase coherence (ITC), as

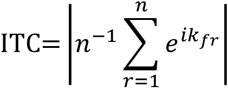

where *n* is the number of trials, and *k* is the phase angle on trial *r* at the frequency *f*. ITC reflects the degree to which the phase angle of an oscillation at any given time relative to a stimulus is consistent across trials. ITC ranges between 0 and 1, with zero indicating uniformly distributed phase angles, and 1 indicating identical phase angles across trials. Statistical comparisons between conditions were carried out using paired *t*-tests in humans, and independent samples *t*-tests in monkeys.

#### Task performance

To analyze accuracy and reaction times of the 1-back and RSVP tasks, we first excluded trials with outliers in the reaction times per subject using the estimator *Sn* [35], at a threshold of 2, as well as trials in which reaction time exceeded 2.5 s. Average accuracy and reaction times per condition (exposed, foil, novel) were then compared by means of planned paired *t*-tests. Figure 4B shows the respective means and corresponding 95% Bayes-bootstrapped (1000 samples) high density intervals from the RSVP task and was plotted using the Robust Statistical Toolbox (https://github.com/CPernet/Robust_Statistical_Toolbox) in Matlab.

#### Pupil-task correlations

To assess the relationship between pupil entrainment during the exposure phase and accuracy in the subsequent test phase, we correlated accuracy in the RSVP task with the normalized spectral power at the 1 Hz pair frequency in the random and structured condition, respectively, using Pearson’s correlation coefficient. To this end, we first normalized the 1 Hz peak by subtracting the average spectral power of four surrounding frequency bins (two above, two below) within each condition. This estimates the distinctiveness of the peak at 1 Hz relative to the surrounding spectrum. To assess the specificity of the correlation between 1 Hz spectral power in the structured condition and accuracy, we further performed partial correlation analyses, controlling for any correlation between 1 Hz spectral power in the random condition and accuracy, again using Pearson’s correlation coefficient. Analyses of (partial) correlations using Spearman’s correlation coefficient yielded the same pattern of results.

#### Effect sizes

For all *t*-tests, we computed Hedges’ *g* and for all ANOVA partial *η*^*2*^ as measures of effect size using the Matlab toolbox Measures of Effect Size (v 1.6.1; https://github.com/hhentschke/measures-of-effect-size-toolbox/) [36].

