## Supplemental Figures for "Pupil diameter tracks statistical structure in the environment"

**Supplemental Information**

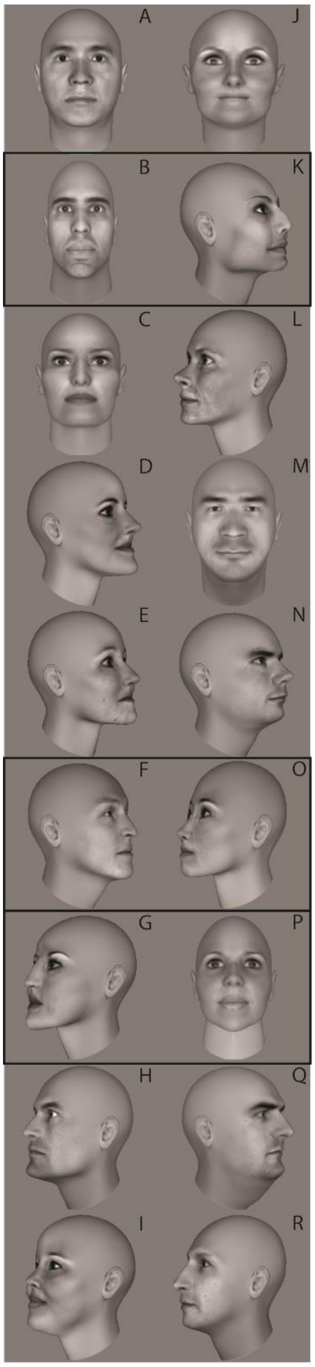


**Fig. S1.** *Face pairs.* Each face image was uniquely paired with one other face image during the exposure phase. Across images, head orientation was balanced such that each of the three head orientations (0°, 60°, 300°) would be followed by each head orientation with equal likelihood. Human subjects were exposed to nine pairs, monkey subjects to three (BK, FO, GP; black boxes).

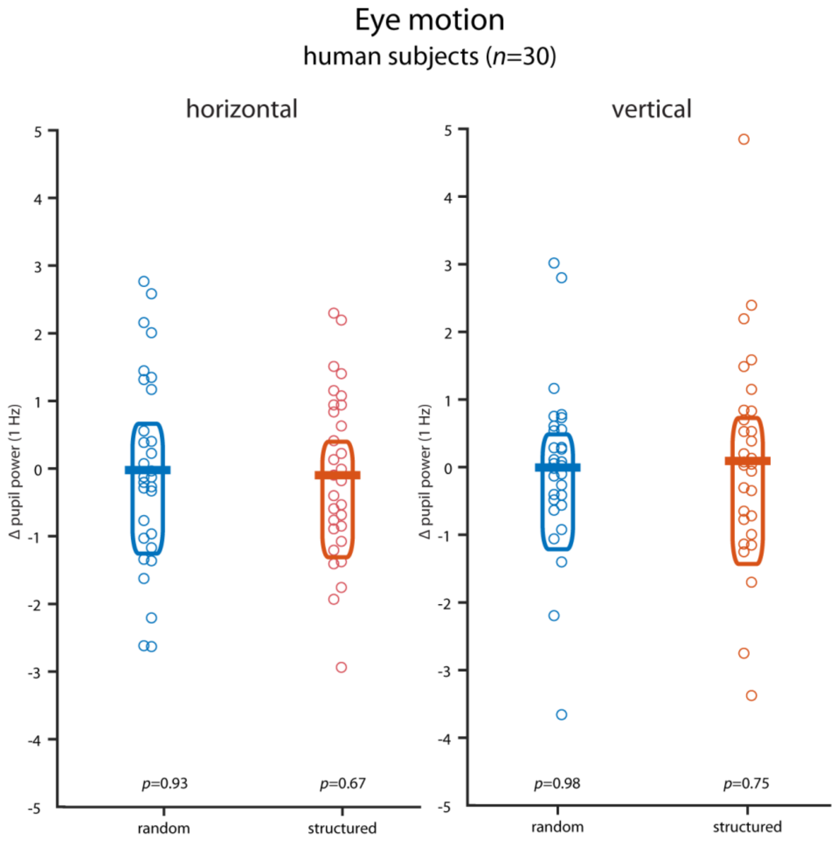


**Fig. S2.** *Eye motion in humans.* There were no statistically significant peaks at 1 Hz in the dynamics of eye position in the random or structured condition in human subjects ($n$=30) relative to the four surrounding frequency bins. Horizontal bars indicate the mean, boxes the 95% Bayes-bootstrapped high density interval, circles the individual subjects' data points, $p$-values are from paired, two-sided $t$-tests of 1 Hz power against the average of four surrounding frequency bins (all $p$>0.6).

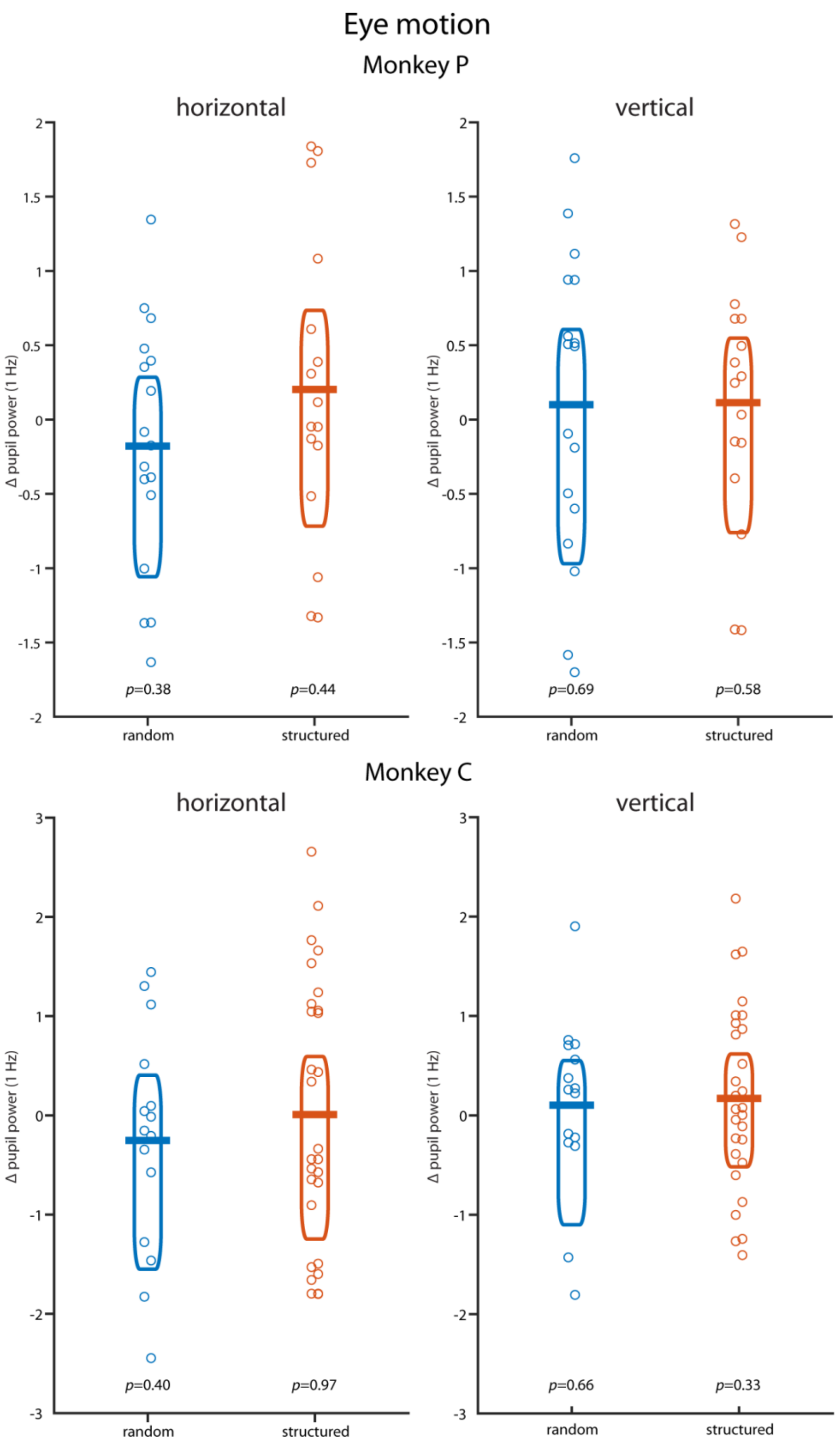


**Fig. S3.** *Eye motion in monkeys.* There were no statistically significant peaks at 1 Hz in the dynamics of eye position in the random or structured condition in monkey P and monkey C relative to the four surrounding frequency bins. Horizontal bars indicate the mean, boxes the 95% Bayes-bootstrapped high density interval, circles the individual runs' data points, *p*-values are from paired, two-sided *t*-tests of 1 Hz power against the average of four surrounding frequency bins (all $p>0.3$).
